# Independent regulation of memory and emotion by selective retrieval

**DOI:** 10.1101/2025.01.02.631155

**Authors:** Yingying Wang, Zijian Zhu, S. Subbulakshmi, Yao Li, Fang Fang, Michael C. Anderson

**Affiliations:** Department of Psychology and Behavioral Sciences, Zhejiang University, Hangzhou, China; School of Psychology, Shaanxi Normal University, Xi’an, China; Department of Psychology, Stanford University, Stanford, USA; Academy for Advanced Interdisciplinary Studies, Peking University, Beijing, China; School of Psychological and Cognitive Sciences and Beijing Key Laboratory of Behavior and Mental Health, Peking University, Beijing, China; Key Laboratory of Machine Perception (Ministry of Education), Peking University, Beijing, China; IDG/McGovern Institute for Brain Research, Peking University, Beijing, China; MRC Cognition and Brain Sciences Unit, University of Cambridge, Cambridge, UK

## Abstract

Selective retrieval of a target memory often triggers inhibitory control to reduce competition from related traces, a process that induces forgetting of the competing content. It is unknown, however, whether inhibition during selective retrieval specifically targets a competitor’s episodic representation or instead extends to its affective components. Here we report evidence in humans that selective retrieval of a neutral memory not only diminishes access to the mnemonic content of unpleasant competing memories, but also alters their emotional character. Memory and emotion suppression effects were accompanied by reduced neural activation and weakened representational patterns unique to competing memories in the VTC and amygdala respectively, in an independent fashion. Selective retrieval engaged the left VLPFC, which decreased in activity over repeated retrievals of the same memory, as competition from the unpleasant scene was resolved. This left VLPFC region colocalizes with key prefrontal regions engaged during cognitive reappraisal, suggesting that selective retrieval’s impact on affective responses may contribute to reappraisal’s benefits. These findings indicate that inhibitory control during selective retrieval affects both mnemonic and affective representations, providing a novel mechanistic basis for a well-known emotion regulation practice.

## Introduction

Re-interpreting an unpleasant memory in a benign way is an effective approach to reducing the negative affect that the memory generates. This emotion regulation strategy, known as cognitive reappraisal (Gross & John, 2003), requires a selective focus on those details that are consistent with a positive reframing that changes the event’s meaning. Such reframing focuses people’s attention on positive interpretations and alters the memory trace so that future retrievals elicit those interpretations (Speer et al., 2021). Although the benefits of such reframing are clear, which element of the refocusing process causes affect regulation effects is less obvious. On the one hand, the ready availability of a positive interpretation may facilitate positive affect whenever an event is remembered; on the other hand, reappraisal might reduce access to unpleasant episodic details and affective content that induce distress. For example, selectively retrieving features needed for a positive interpretation may induce forgetting of all event details that get omitted, via inhibitory mechanisms that suppress competing traces. Selectively retrieving target memories, given a cue, is widely known to induce episodic forgetting of competitors, a phenomenon known as retrieval-induced forgetting (RIF) (Anderson & Hulbert, 2021; Anderson et al., 1994; Marsh & Anderson, 2024; Meyer & Benoit, 2022; Storm & Levy, 2012). Critically, the selective retrieval process that induces RIF engages areas within the left ventrolateral prefrontal cortex (VLPFC) resembling those involved in cognitive reappraisal (Buhle et al., 2014; Yang et al., 2021). These functional and neural parallels raise the hypothesis that reappraisal benefits derive, in part, from memory inhibition (Engen & Anderson, 2018). Here we test this possibility, which we refer to as the selective retrieval hypothesis of cognitive reappraisal.

If reappraising an unpleasant event induces RIF for omitted content, inhibitory processes may reduce negative affect in one of two broad ways. On the one hand, given that the details of episodic memories may generate emotional states (Engen & Anderson, 2018; Morina et al., 2011), selective retrieval of benign event details may reduce negative affect indirectly by inhibiting event features that might otherwise trigger an unpleasant emotional response (Marche et al., 2016; Wauters et al., 2024). In such cases, any tendency for selective retrieval to regulate emotion would depend directly on the amount of episodic retrieval-induced forgetting. On the other hand, selective retrieval may regulate emotion directly by inhibiting emotional learning associated to the competing event details. Such affective inhibition may occur in parallel to inhibition affecting episodic representations of the competing traces. Precedent for such a parallel regulation of episodic and affective representations derives from research on intentional retrieval suppression, which has shown that suppressing the retrieval of unpleasant scenes evokes parallel inhibitory control by the right DLPFC over both the hippocampus and the amygdala (Anderson et al., 2024; Gagnepain et al., 2017). If affective and mnemonic inhibition occur in parallel, any affect reduction induced by selective retrieval need not be related to the amount of RIF people exhibit. Moreover, whereas both the indirect and direct accounts predict reduced activity in emotion-related regions, only the direct account posits that inhibition acts directly on emotion-related traces.

Importantly, little evidence exists about whether selective retrieval diminishes affective responding to unpleasant competing memories. To address whether RIF induces affective inhibition, we tested whether selective retrieval of neutral target memories associated to a cue altered the emotional tone of competing stimuli and whether this effect arises as a specific consequence of retrieval-related inhibitory processes. We trained participants to associate two unrelated stimuli, a neutral word and a negative picture, to a common retrieval cue. Participants then performed selective retrieval practice on the neutral word using the common cue without retrieving the affective picture. If RIF extends to emotional responses, retrieving a neutral word should diminish both the episodic recall of the competing unpleasant picture as well as its emotional representations, even though the word being retrieved has nothing at all to do with re-interpreting the scene. If RIF constitutes a fundamental mechanism for reappraisal, as suggested by the selective retrieval hypothesis, selective retrieval should engage brain regions overlapping with the key brain areas involved in cognitive reappraisal. Furthermore, we examined whether any emotion regulation induced by selective retrieval arose by impairing episodic memory for scene details or by direct inhibition of affective representations supported by the amygdala.

## Results

### Retrieval induces forgetting of unpleasant competing memories

We first examined whether retrieving a neutral memory impaired the retention of the unrelated unpleasant competitor scene in a behavioral experiment (Exp. 1, Fig. 1A). We calculated the percentage of targets and competitors that were recalled on the final test (Fig. 1B) separately for the Retrieval, Restudy, and Control conditions. A repeated measures ANOVA revealed a significant main effect on competitor recall accuracy across the three conditions (F(2,70) = 6.13, p = .004, *η*_p_^2^ = 0.15). Replicating RIF, selectively retrieving neutral words, given the shared cue, robustly impaired later recall of the competing scenes. Specifically, recall accuracy for competing unpleasant scenes decreased in the Retrieval condition compared to the recall of competing scenes in the Control condition (t(35) = -2.74, p = .010, Cohen’s d = -0.46) and also when compared to the recall of competing scenes within the Restudy condition (t(35) = -3.42, p = .002, Cohen’s d = -0.57). In contrast, recall accuracy for unpleasant scenes in the Restudy condition did not differ from that observed in the Control condition (t(35) = 0.17, p = .865, Cohen’s d = 0.03), confirming the widely observed finding that RIF is retrieval-specific. Unsurprisingly, target recall also varied significantly across the three conditions (F(2,70) = 31.5, p < .001, *η*_p_^2^ = 0.47), with recall improving for items in the Retrieval (t(35) = 5.47, p < .001, Cohen’s d = 0.91) and Restudy conditions (t(35) = 6.54, p < .001, Cohen’s d = 1.09) relative to recall observed for Control items. Indeed, Restudying targets improved their recall even more than did Retrieval practice (t(35) = 2.88, p = .007, Cohen’s d = 0.48). These findings support the hypothesis that impaired recall of competitors in the Retrieval condition derives from processes specific to retrieval and not simply from strengthening target items.

**Figure 1.**
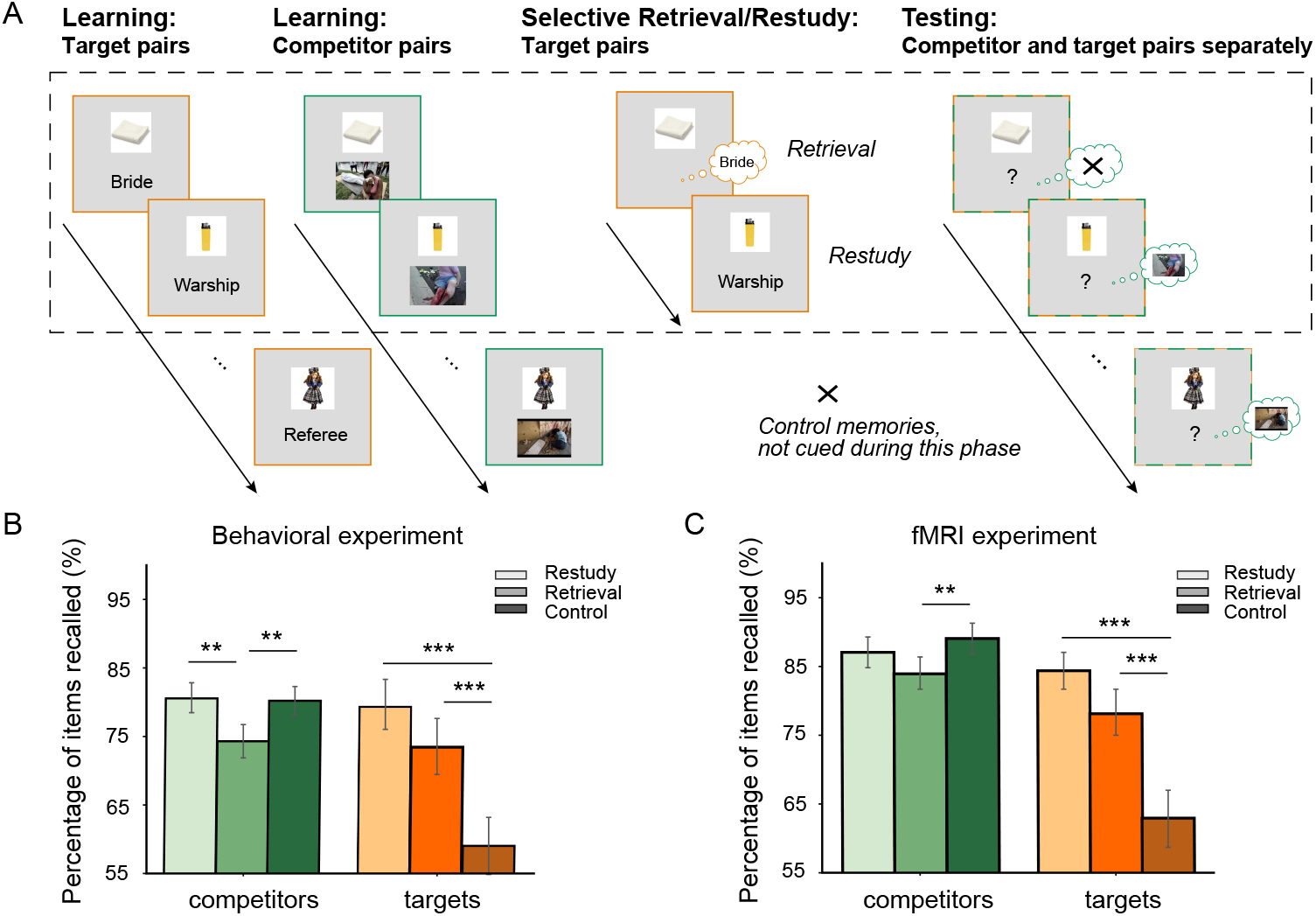
Schematic of the experimental procedure and the behavioral results. **(A)** The procedure for Experiments 1 and 2 (excluding the pattern localizer and emotional response tests in Experiment 2). We trained participants on object-word (target) and object-scene (competitor) pairs in separate blocks. Afterwards, participants performed selective Retrieval/Restudy on only object-word associates. During Retrieval trials, we cued participants with an object (eight times each across the entire phase) and asked them to retrieve the associated word (the target), with the unpleasant scene (the competitor) assumed to interfere. During Restudy trials, we presented participants with a complete object-word associate (eight times each across the entire phase) and asked them to silently repeat the associate. We omitted some of the studied pairs during this phase so they served as a Control against which we could assess the impact of selective Retrieval/Restudy on both targets and competitors. Participants completed this Selective Retrieval/Restudy phase inside the fMRI scanner in Experiment 2. In the testing phase, we presented object cues for participants to retrieve the associated scenes or words. We always tested recall for the scenes first, followed by a separate testing block for the words. The colored frames illustrate the item types and were not visible to participants. **(B)** Behavioral results from the final recall test in Experiment 1. Retrieval and Restudy both improved recall performance for target words; however, only Retrieval and not Restudy on target words induced forgetting of the competing scenes. **(C)** Behavioral results from the recall test in Experiment 2, replicating the retrieval-induced forgetting found in Experiment 1.** p < .01, *** p < .001. (Two-tailed t test). Error bars indicate the standard errors of the mean.

We replicated the foregoing behavioral patterns in an fMRI experiment (Exp. 2, Fig. 1C) conducted with an independent set of participants. Again, selective Retrieval on neutral memories caused significant RIF on competing unpleasant scenes (t(27) = -3.02, p = .006, Cohen’s d = -0.57), but Restudy did not (t(27) = -0.91, p = .370, Cohen’s d = -0.17). Both Retrieval (t(27) = 4.60, p < .001, Cohen’s d = 0.87) and Restudy (t(27) = 6.34, p < .001, Cohen’s d = 1.20) improved later retention for the neutral targets significantly. Taken together, these attributes show that active retrieval of a memory disrupts later retention of unrelated competing unpleasant scenes. Critically, this memory disruption occurred even though the retrieved word itself was neutral in valence and even though retrieval practice involved no reinterpretation or reappraisal of the competing memory. Thus, simply accessing neutral content associated to a shared cue is sufficient to impair competing memories, even if those competitors are unpleasant, independent of whether reinterpretation occurs. This finding reinforces the possibility that the positive reinterpretations generated during reappraisal are not necessarily the primary cause of the emotional benefits of reappraisal, and instead point to selective retrieval of content in support of those reinterpretations.

### Retrieval reduces neural responses representing competing memories

If selective retrieval induces RIF because it engages inhibitory control to suppress competitors, retrieval should suppress scene-related activity in brain structures involved in episodic representation and reinstatement, such as the hippocampus and the ventral temporal cortex. Therefore, by our inhibition hypothesis, Retrieval should suppress activity relating to the reinstatement of the competing scene, whereas Restudy should not. Given this difference, our hypothesis predicts diminished overall activity and diminished competitor reinstatement during Retrieval compared to Restudy. If such differences occur and are caused by inhibition, both Retrieval activity and reinstatement should predict later RIF, such that larger reductions during Retrieval are linked to greater forgetting.

To test these predictions, we performed a wholebrain analysis comparing Retrieval versus Restudy, which showed significantly decreased activation in the left HPC and VTC during Retrieval compared to Restudy (Fig. 2A; Table 1). An ROI analysis confirmed significant activation decreases in the Retrieval condition relative to the Restudy condition in the left HPC (Fig. 2B, t(27) = -4.39, p < .001, Cohen’s d = 0.83) and VTC (Fig. 2D, t(27) = -10.92, p < .001, Cohen’s d = 2.06). To test whether this decreased activity was related to RIF, we performed a Pearson correlation between the ROI averaged activation and the forgetting effect. Whereas the HPC is critical for memory, we observed no reliable correlations between left HPC activation and forgetting effects in either the Retrieval (Fig. 2C, r(26) = -0.16, p = 0.401) or the Restudy (Fig. 2C, r(26) = 0.30, p = 0.124) condition. In contrast, the correlation between activation in VTC and RIF was significant, with lower activation during selective Retrieval predicting greater RIF (Fig. 2E, r(26) = -0.41, p = 0.033); the same correlation was not significant in the Restudy condition (Fig. 2E, r(26) = -0.18, p = 0.351). These findings suggest that activity reductions during selective retrieval may have been involved in inducing later forgetting, at least in the VTC.

**Table 1.** Regions showing a difference in activity between Retrieval and Restudy trials ^a^.

| Anatomical description | No. of voxels | MNI coordinates in mm |  |  | Peak z |
| --- | --- | --- | --- | --- | --- |
|  |  | x | y | z |  |
| Retrieval > Restudy |  |  |  |  |  |
| Left middle/inferior frontal gyrus | 247 | -50 | 22 | 28 | 4.30 |
| Left frontal pole | 186 | -28 | 48 | 7 | 3.89 |
| Right Frontal pole | 157 | 22 | 62 | 7 | 3.45 |
| Paracingulate gyrus (anterior) | 1246 | 0 | 14 | 51 | 5.41 |
| Posterior cingulate gyrus | 236 | -2 | -40 | 23 | 4.10 |
| Left insula | 648 | -34 | 24 | 2 | 5.25 |
| Right insula | 526 | 33 | 26 | -6 | 5.34 |
| Left precuneus | 232 | -12 | -72 | 32 | 5.32 |
| Right precuneus | 87 | 10 | -74 | 39 | 4.21 |
| Occipital pole | 769 | 14 | -104 | 7 | 5.10 |
| Retrieval < Restudy |  |  |  |  |  |
| Left hippocampus | 338 | -22 | -26 | -9 | 4.15 |
| Left amygdala | Same cluster | -18 | -8 | -14 | 3.86 |
| Medial frontal cortex | 1640 | 2 | 56 | -11 | 4.51 |
| Right middle frontal gyrus | 1274 | 34 | 38 | 44 | 4.23 |
| Left middle frontal gyrus | 767 | 28 | -4 | 49 | 4.37 |
| Left supramarginal gyrus | 336 | -64 | -36 | 32 | 4.28 |
| Right supramarginal gyrus | 111 | 60 | -30 | 42 | 4.20 |
| Occipital parietal cortex | 21905 | -39 | -78 | -17 | 6.42 |

<sup>a</sup>Peak coordinates of the regions showing greater activity for Retrieval relative to Restudy items (and vice versa) at a cluster-defining threshold of $z > 2.3$ ( $P < 0.01$ ) and a cluster probability of $P_{\text{corr}} < 0.05$ corrected for multiple comparisons.

**Figure 2.**
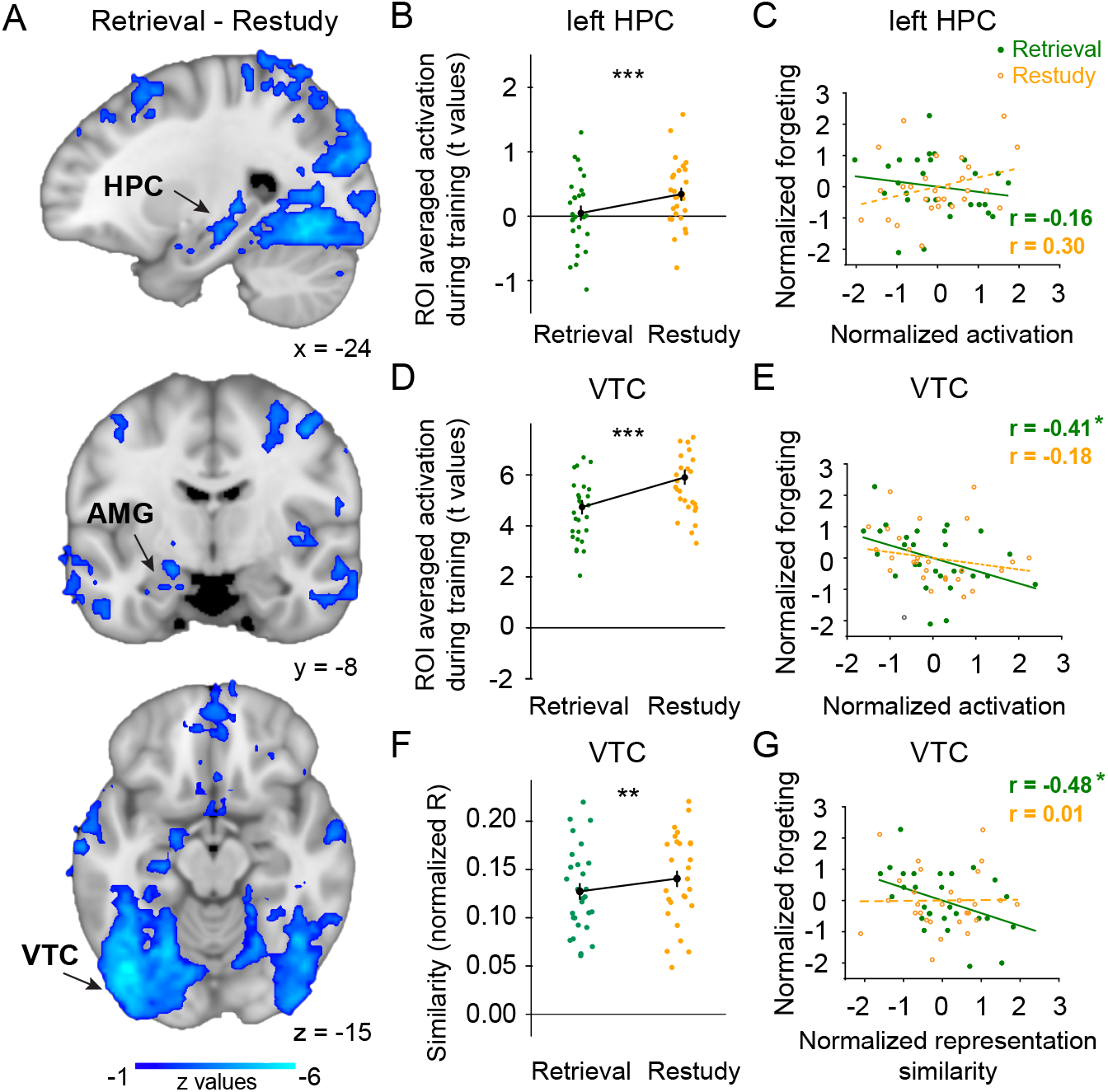
Impact of target retrieval on competing memories in HPC and VTC. **(A)** Voxels showing decreased activation in the left HPC, left AMG, and VTC during Retrieval compared to during Restudy. The color bar indicates z values (corrected). **(B)** Mean univariate activation during Retrieval was significantly lower than that during Restudy in the left HPC. **(C)** Left HPC activation did not predict the forgetting effect in either the Retrieval (solid line, r(26) = -0.16, p = 0.401) or Restudy (dotted line, r(26) = 0.30, p = 0.124) condition. **(D)** Mean univariate activation during Retrieval was significantly lower than that during Restudy in VTC. **(E)** VTC activation was significantly correlated with the forgetting effect in the Retrieval condition (solid line, r(26) = -0.41, p = 0.033) but not in the Restudy condition (dotted line, r(26) = -0.18, p = 0.351). Lower VTC activation during Retrieval predicted a larger RIF effect. **(F)** The representational similarity to scene templates was lower during Retrieval than during Restudy in VTC. **(G)** The representational similarity in VTC significantly correlated with the forgetting effect in the Retrieval (solid line, r(26) = -0.48, p = 0.010) but not in the Restudy (dotted line, r(26) = 0.01, p = 0.948) condition. Weaker competitor activation in VTC during Retrieval, but not Restudy predicted a larger RIF effect. For panels B, D, and F, ** p < .01, *** p < .001, (Two-tailed t test). Error bars indicate the standard errors of the mean. For panels E and G, * p < .05, (Two-tailed Pearson correlation).

To test whether similar effects arose in neural pattern representations, we developed an index sensitive to the neural representations of each competitor by using a template-based pattern-tracking approach (Wimber et al., 2015). We repeatedly exposed participants to each competing picture in an independent picture-viewing procedure as they were scanned with fMRI. For each picture, we obtained activation-pattern templates, representing the canonical response to viewing the picture. We then used these templates in a representational similarity analysis (RSA) in which we estimated how much, during Retrieval/Restudy, those canonical patterns for competitors arose (see Fig. 5A). This procedure thus estimates how much each competing scene was reactivated during Retrieval or Restudy of target items. During Retrieval trials, we expected that inhibition would reduce competitor reactivation, rendering the remaining neural pattern less similar to its template. We thus performed RSA between activation patterns during each Retrieval/Restudy trial with its competitor’s template (within-item RSA) and with the activation pattern for a random other item (across-item RSA). We found decreased representational similarity for competing scenes (within-item RSA) during the Retrieval condition compared to the Restudy condition in VTC (Fig. 2F, t(27) = -3.33, p = .003, Cohen’s d = -0.63). This finding suggests reduced reinstatement of competitor patterns during selective Retrieval, consistent with an impact of inhibitory control on pattern representations. However, the within-item RSA during the Retrieval condition was not decreased relative to the Restudy condition in the left HPC (t(27) = -0.34, p = .738, Cohen’s d = -0.06). Nevertheless, the level of representational similarity we observed for competitors in VTC predicted later forgetting in the Retrieval condition (Fig. 2G, r(26) = -0.48, p = 0.010) but not in the Restudy condition (r(26) = 0.01, p = 0.948) condition, such that lower competitor reactivation was linked to a greater RIF effect. For the left HPC, the level of representational similarity for competitors did not predict later forgetting in the Retrieval condition (r(26) = -0.22, p = 0.263), but positively predicted later forgetting in the Restudy condition (r(26) = 0.47, p = 0.011). These findings suggest, as was found in Wimber et al. (2015), that memory representations in VTC for competing scenes were suppressed during selective retrieval and that this suppression contributed to RIF for those scenes.

### Repeated retrieval reduces prefrontal control over memory representations

We next examined whether reduced reactivation for competitors in VTC was linked to prefrontal markers of the successful resolution of competition. Multiple lines of work using fMRI, EEG, pupillometry and even cFOS imaging in rodents indicates that the prefrontal cortex grows less engaged over repeated selective retrieval trials as interference from competing memories is gradually resolved (Anderson & Hulbert, 2021). This reduced need for metabolically costly control processes is viewed as a benefit of resolving competition and has been linked to (a) the successful forgetting of competing memories (Kuhl et al., 2007; Wimber et al., 2015), and (b) reduced cortical pattern reinstatement for competitors over repetitions (Wimber et al., 2015). If reductions in item-specific pattern evidence during selective retrieval derive from the engagement of prefrontal control, the current findings should replicate the conflict reduction benefit in left VLPFC and link it to VTC pattern suppression.

To test this hypothesis, we extracted the voxels in the left middle/inferior frontal gyrus (M/IFG) and first compared the averaged activation within the region between the Retrieval and Restudy conditions. This ROI-level analysis confirmed significantly higher activation in the left M/IFG (hereinafter, VLPFC) during Retrieval than during Restudy (Fig. 3A, t(27) = 2.18, p = .038, Cohen’s d = 0.41). To establish the conflict reduction benefit, we examined how this left VLPFC activation changed over quartiles (Fig. 3B). A two (conditions: Retrieval vs. Restudy) by four (quartiles) ANOVA showed a significant interaction (F(3,78) = 6.82, p < .001, *η*_p_^2^ = 0.21). Replicating previous work (Kuhl et al., 2007), left VLPFC activation decreased over quartiles (main effect of Retrieval repetition quartiles: F(3,78) = 5.74, p = .001, *η*_p_^2^ = 0.18) with greater activation in the first than the remaining quartiles (1st vs. 2nd: t(27) = 3.97, p = .001, Cohen’s d = 0.75; 1st vs. 3rd: t(27) = 4.16, p < .001, Cohen’s d = 0.79; 1st vs. 4th: t(27) = 3.55, p = .005, Cohen’s d = 0.68). A modest reduction was found in the Restudy condition (main effect of Restudy repetition quartiles: F(3,78) = 2.45, p = .070, *η*_p_^2^ = 0.09). Thus, as in prior work, we observed that demands on prefrontal control declined over retrieval practice trials consistent with prior work on the conflict reduction benefit (Anderson & Hulbert, 2021).

**Figure 3.**
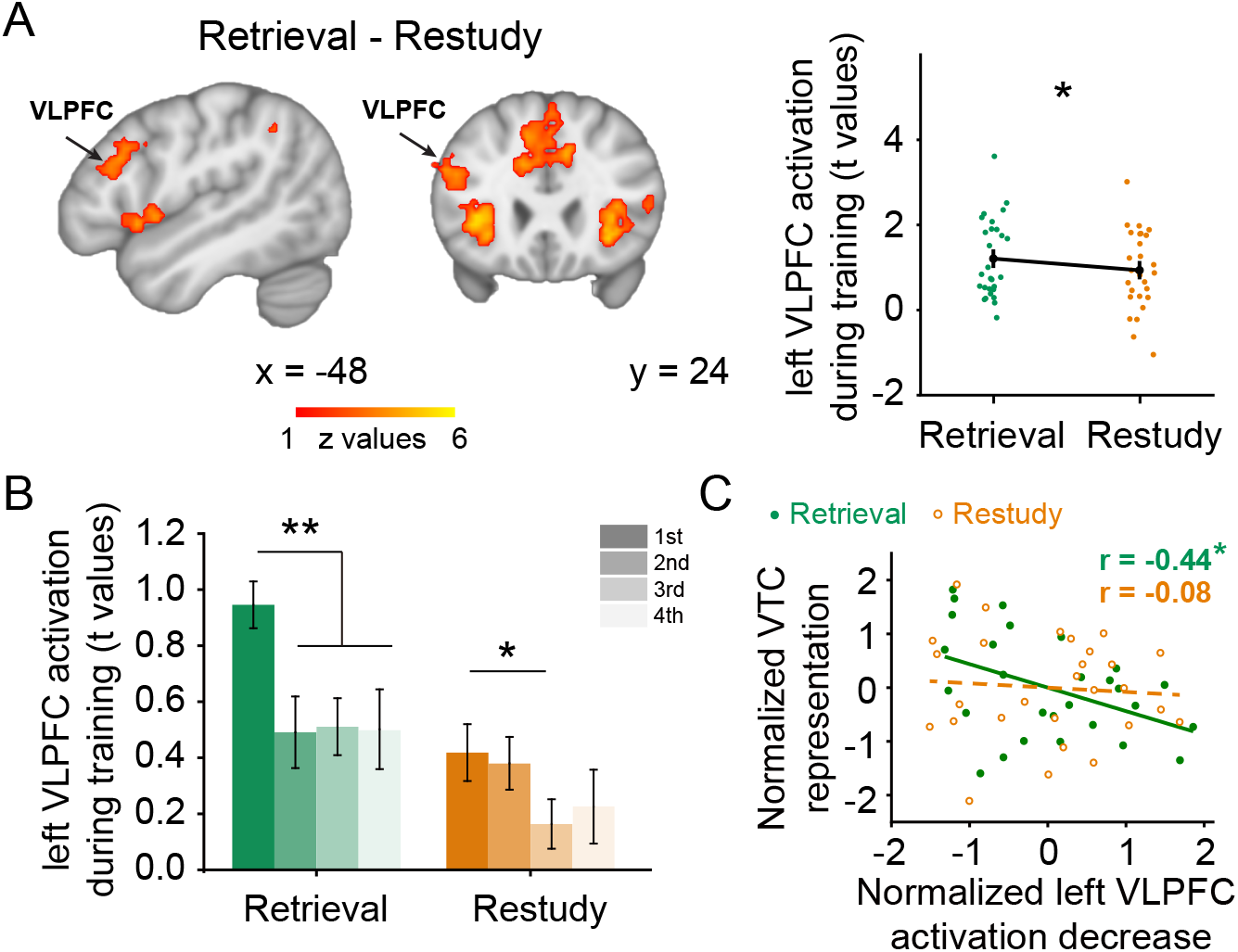
Activation of the left VLPFC in selective Retrieval. **(A)** (left) Voxels regions showing greater activation in the left VLPFC during Retrieval than during Restudy. The color bar indicates z values (corrected). (right) Activation in the left VLPFC showed greater activation during Retrieval than during Restudy. **(B)** Activation in left VLPFC ROI decreased along with repeated practice in the Retrieval condition. * p < .05, ** p < .01 (Two-tailed t test). Error bars indicate the standard errors of the mean. **(C)** The activation decrease in the left VLPFC predicted representational similarity in VTC during selective Retrieval (solid line, r(26) = -0.44, p = 0.020). No such correlation was found in the Restudy condition (dotted line, r(26) = -0.08, p = 0.685). * p < .05, (Pearson correlation).

Having established a robust conflict reduction benefit, we next sought to determine whether the magnitude of this benefit was related to disruptions in item-specific representations underlying the competitor. To achieve this, we quantified, for each subject, the extent to which VLPFC activation declined over blocks and related this to the magnitude of the representational disruption. Consistent with prior work, we observed that larger conflict reduction benefits in the Retrieval condition were associated with lower representational similarity for competitors in the VTC (Fig. 3C, r(26) = -0.44, p = 0.020). No such correlation was found in the Restudy condition (Fig. 3C, r(26) = -0.08, p = 0.685). These findings link reductions in reinstatement for unpleasant scenes in VTC during selective retrieval to the putative engagement of inhibitory control mechanisms in left VLPFC.

### Retrieval reduces affective responding to competing memories

Having established the detrimental effect of selective retrieval on behavioral and neural indices of episodic memory our central aim was to determine whether selective retrieval’s impact extended to brain regions involved in negative affect such as the Amygdala (AMG). Notably, such an impact would be surprising, given that our selective retrieval practice task made no reference to the need to regulate emotion, and in no way required re-appraisal of an unpleasant stimulus, yet such an impact is predicted by our main hypothesis. Specifically, we predicted that AMG activity should be reduced during Retrieval trials, in which inhibitory control is predicted to be engaged to inhibit unpleasant competitor memories, compared to Restudy trials, in which these processes are not engaged. Consistent with this hypothesis, our whole-brain analysis comparing Retrieval versus Restudy conditions indeed revealed significantly decreased activation in the left AMG during Retrieval compared to Restudy (Fig. 2A; Table 1). An ROI analysis further confirmed an activation decrease in the Retrieval condition compared to the Restudy condition in the left AMG (Fig. 4A, t(27) = -4.54, p < .001, Cohen’s d = 0.86). Interestingly, left AMG activation was significantly lower during Retrieval trials than it was in the fixation baseline (Fig. 4A, t(27) = -1.98, p = .029, Cohen’s d = 0.37, one tailed), suggesting that lower AMG activity does not simply reflect a failure to engage AMG, but rather an active reduction during selective retrieval. Because the AMG was unlikely to be affected by retrieving affectively neutral target words, below-baseline activation in the left AMG likely reflects processes engaged to suppress affective responses to the competing scene. Taken together, these findings provide evidence consistent with the possibility that inhibitory control initiated by active retrieval suppresses affective representations related to the unpleasant scene.

**Figure 4.**
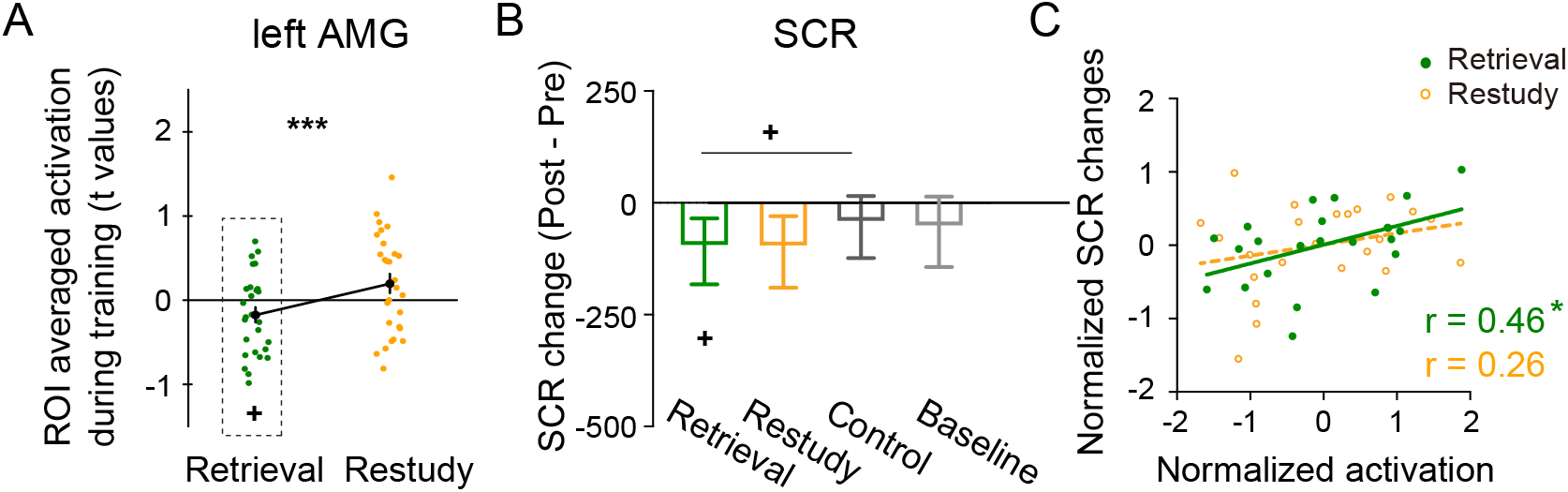
Affective response decreases during Retrieval. **(A)** Mean activation in the ROIs of the left AMG. The left AMG showed below-baseline activation during Retrieval (shown in the dotted frame). *** p < .001, (Two-tailed t test). + p < .05, (One-tailed t test). Error bars indicate the standard errors of the mean. **(B)** SCR changes in the Post- versus Pre-experiment tests for each condition. The SCR decrease in the Retrieval condition was larger than that in the control condition. + p < .05, (One-tailed Wilcoxon test). Error bars indicate the standard errors of the mean. **(C)** Correlations between the neural activation and SCR effects. Left AMG activation was correlated with the SCR changes in the Retrieval (r(21) = 0.46, p = 0.027) but not in the Restudy (r(21) = 0.26, p = 0.232) condition. Lower activation in left AMG during Retrieval predicted larger SCR decreases for competing items after retrieval practice, compared to before. * p < .05, (Pearson correlation).

Reductions in amygdala activity during selective retrieval do not necessarily establish that selective retrieval regulated affective content of the competing memories, however. To explore this possibility, we examined whether inhibitory processes triggered by selective retrieval durably affected emotional responses to competing scenes. Exp. 2 measured participants’ emotional responses to the competitors before and after Retrieval/Restudy, with two tests: (1) a skin conductance response (SCR) test to objectively measure changes in psychophysiological responses to the unpleasant scene, and (2) a Valence/Arousal rating, to measure changes in subjective valence. If the reduced amygdala activation reflects the impact of control on emotion representations, a durable effect on SCR and subjective measures may arise in the Retrieval condition. Because the SCR measure did not follow a normal distribution, we performed non-parametric statistical analysis with the Wilcoxon signed-rank test on this measure. Comparing the Post-test vs. Pre-test SCR change indices in the Retrieval/Restudy condition with the analogous changes in the Control condition we indeed found a larger SCR decrease in the Retrieval condition than in the Control condition (Fig. 4B, W = 83, p = .049, r_rb_ = -0.40, one tailed). The comparable SCR changes in the Restudy condition did not differ from those in the Control condition (W = 127, p = .377, r_rb_ = -0.08, one tailed). Moreover, these SCR changes to unpleasant competing scenes (Post- vs Pre-Retrieval) were related to univariate activation in the left AMG during selective retrieval (Fig. 4C, r(21) = 0.46, p = 0.027), consistent with a role of active AMG suppression in altering longer-lasting affective responding. In contrast, no correlation was observed between SCR change and univariate activation in the AMY in the Restudy condition (Fig. 4C, r(21) = 0.26, p = 0.232).

Next, we examined the analogous Post-Test vs Pre-test change indices on our subjective emotional rating measures of Valence and Arousal. Unlike our objective index, these rating changes did not differ across conditions (ps > .50). Therefore, psychophysiological responses, but not subjective ratings of emotion showed evidence of inhibition-related changes. Indeed, subjective and objective emotion indices do not always follow together, with the latter influenced by activity in a broader set of cortical regions than the former, which may be more directly tied to the threat detection processes mediated by the amygdala (Barrett et al., 2007; LeDoux, 2014; LeDoux & Brown, 2017; LeDoux & Pine, 2016; Zhou et al., 2021).

### Retrieval suppressed item-unique neural patterns representing emotional responses

Selective retrieval decreased neural activation in the left AMG and also decreased SCRs on the final test. This pattern suggests that selective retrieval engages inhibitory control to reduce emotional responses to competing scenes and that these reductions induce persisting affective changes. Persisting SCR reductions may be mediated by disruption of item-unique representations within the AMG. To test this possibility, we used the canonical template-tracking method used to analyze our VTC findings to estimate how much, during Retrieval/Restudy, those canonical patterns for competitors arose (Fig. 5A). This procedure thus estimates the state of each competing scene during Retrieval or Restudy of target items (Fig. 5A). If such a reduction in competitor reactivation arose in the amygdala, it would reinforce the possibility that selective retrieval of neutral content can indeed impact the emotional content of competing memories.

We thus performed RSA between activation patterns during each of the Retrieval/Restudy conditions with their corresponding competitor’s template (within-item RSA) and with the activation pattern for a random other item (across-item RSA). Critically, we found that in the Retrieval condition, during retrieval of the neutral target word, the representational similarity of activation patterns in the amygdala to the template for the competing scene was even lower than it was to a random other item template (Fig. 5B left, Withinvs. Across-item RSA: t(27) = -2.50, p = .019, Cohen’s d = -0.47). This finding suggests that item-unique information had indeed been suppressed in the left AMG. In contrast, in the Restudy condition, no withinvs. across-item representational similarity differences arose in the left AMG (Fig. 5B right, Withinvs. Across-item RSA: t(27) = -0.35, p = .728, Cohen’s d = 0.07). This below-baseline representational similarity to competitor templates supports the possibility that in-hibitory control processes may have acted on the amygdala to disrupt the retention of item-unique information.

**Figure 5.**
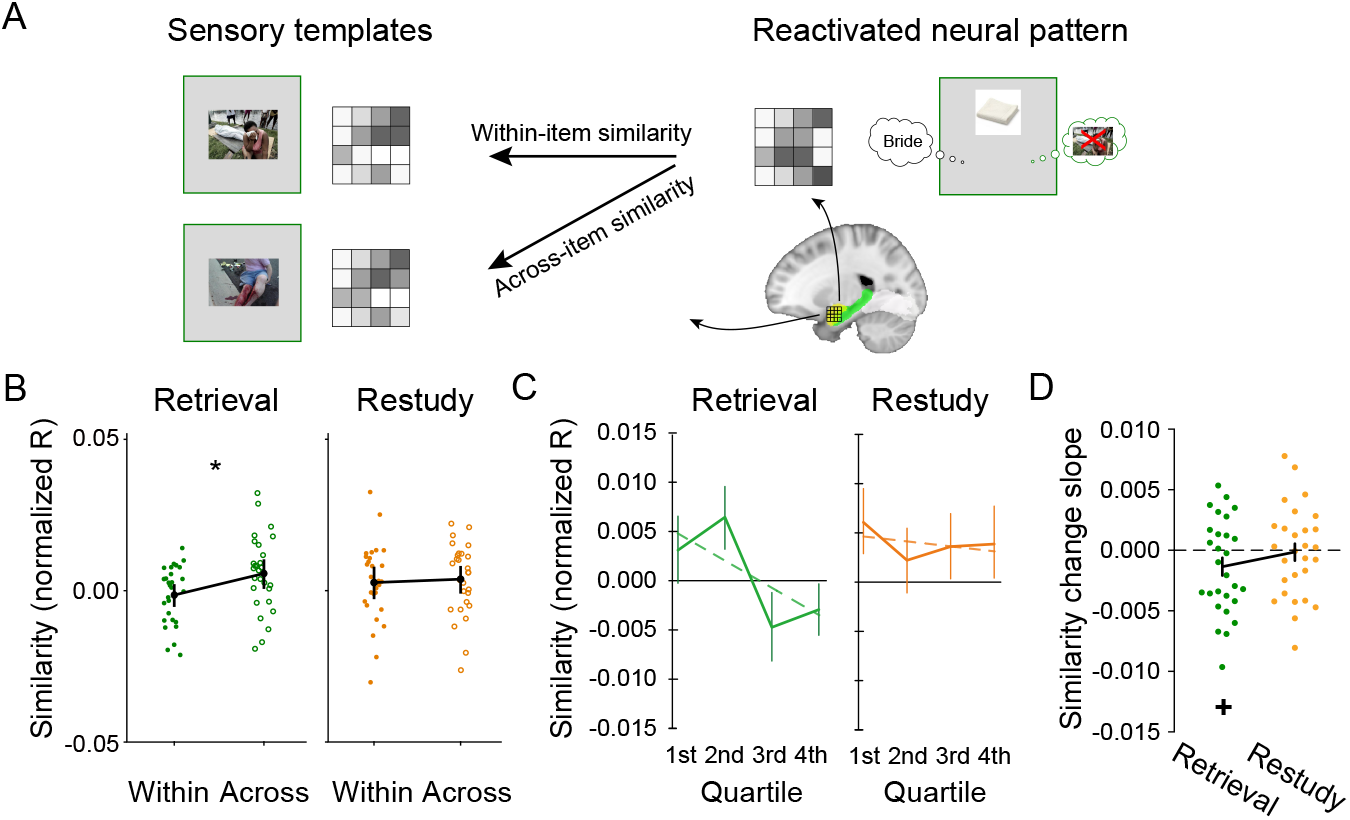
Reduced representational similarity in AMG. **(A)** Our representational similarity analysis approach. Pearson correlation was performed between activation patterns during Retrieval/Restudy in each ROI and those patterns elicited by viewing the corresponding competitor picture. **(B)** Within-item representational similarity was lower than across-item similarity during Retrieval in the left AMG. * p < .05, (Two-tailed t test). Error bars indicate the standard errors of the mean. **(C)** A gradual representational similarity decrease was observed for Retrieval quartiles. No corresponding decrease was observed for Restudy trials. The dotted line represents a linear fit for similarity across the four quartiles. **(D)** The mean slope of representational similarity decline using a linear fit across Retrieval/Restudy quartiles. + p < .05, (One-tailed t test).

The foregoing findings, however, do not provide insight into the dynamics of how such below-baseline pattern similarity might develop over the course of the experiment. For example, in prior selective retrieval research, the match of the competing memory’s template to the activation patterns observed during a retrieval practice trial declined progressively over repetitions, as the competing item was gradually suppressed (Wimber et al., 2015). If an analogous process is at work in the modulation of amygdala representations, we should find a corresponding decline in pattern similarity over repetitions. To test this possibility, we combined voxels from the bilateral amygdala and computed RSA for each Retrieval/Restudy quartile during the retrieval practice phase. Indeed, we observed gradually reduced competitor reactivation in bilateral AMG for the Retrieval condition (Fig. 5C left). To quantify the rate at which competitor reactivation declined, we calculated the slope of representational similarity decline using a linear fit across Retrieval/Restudy quartiles (dotted lines in Fig. 5C). The mean slope in the Retrieval condition was significantly lower than 0 (Fig. 5D, t(27) = -1.81, p = .041, Cohen’s d = 0.34, one tailed). In contrast, no competitor reactivation decrease was evident during Restudy trials (Fig. 5D, independent t-test on the mean linear fit slope: t(27) = -0.22, p = .412, Cohen’s d = 0.04, one tailed). These findings suggest that retrieving an entirely neutral and unrelated target not only reduced amygdala activation, but also may disrupt representational patterns associated with unpleasant competing memories, and that this disruption builds progressively over selective retrieval attempts.

### Emotional response decreases are not related to memory decreases

Decreased emotional responding to unpleasant scenes may be driven directly by inhibition acting on representations in AMG, indirectly by reduced memory for the unpleasant scene, or by both direct and indirect influences. The foregoing evidence that changes in SCR are related to reduced activity in AMG support direct inhibition as one factor. But might altered affective responding also be related to RIF in episodic memory? To test whether decreased emotional responding depended on the forgetting effect, we correlated reduced activity in the AMG with RIF. We found that amygdala activity (Retrieval: r(26) = 0.15, p = 0.461; Restudy: r(26) = -0.16, p = 0.416) was not associated to forgetting due to Retrieval (i.e., Retrieval – Control) or Restudy (i.e., Restudy - Control). These findings are echoed in the relationships of SCR to RIF. Pearson correlation analysis showed that emotional changes on the SCR test (i.e., the SCR decrease for competing scenes) was uncorrelated with memory decreases due either to the Retrieval (r(21) = 0.24, p = 0.268) or to the Restudy (r(21) = -0.13, p = 0.544) of target items. Taken together, these findings suggest that reduced affective responding (as reflected either in psychophysiological responses to competing memories or in neural indices) may not have originated from disrupted episodic memory for the competing scene. Converging with this interpretation, neither VTC activation (Retrieval: r(21) = 0.20, p = 0.364; Restudy: r(21) = -0.24, p = 0.275) nor VTC representational similarity (Retrieval: r(21) = 0.07, p = 0.754; Restudy: r(21) = -0.05, p = 0.834) showed relationships to SCR changes. These findings raise the possibility that emotional response decreases induced by selective retrieval may originate from the modulation of distinct neural structures and may be partially independent from those underlying episodic memory deficits observed in RIF.

### Retrieval modulates the functional connection between the left AMG and the VLPFC

If the reduced neural activation that we have observed in the AMG reflects the impact of top-down inhibitory control processes, greater negative coupling with this region should occur during Retrieval compared to Restudy trials, in which inhibitory control should not be engaged. To test this hypothesis, we performed a generalized PPI (gPPI) analysis examining variations in connectivity arising across Retrieval and Restudy conditions between the left amygdala and other regions. The gPPI measures condition-dependent changes in functional connectivity between a seed on a target region after partialing out task-unrelated connectivity and task-related activity. Using the left AMG as a seed, we observed significantly decreased functional connectivity between AMG and the left VLPFC (Fig. 6A) in the Retrieval compared to the Restudy condition (x = -54, y = 24, z = 10, t(27) = -5.57, p < 0.001). Specifically, negative coupling was observed in the Retrieval condition (Fig. 6B). This finding is consistent with the possibility that the left VLPFC negatively modulated activity in the left AMG in the Retrieval condition, although the direction of influence cannot be ascertained from this method alone. Nevertheless, these findings are compatible with the common assumption that left VLPFC contributes to a network that modulates affective activity in the left AMG, supporting the potential role of selective retrieval as a mechanism contributing to the benefits of reappraisal.

**Figure 6.**
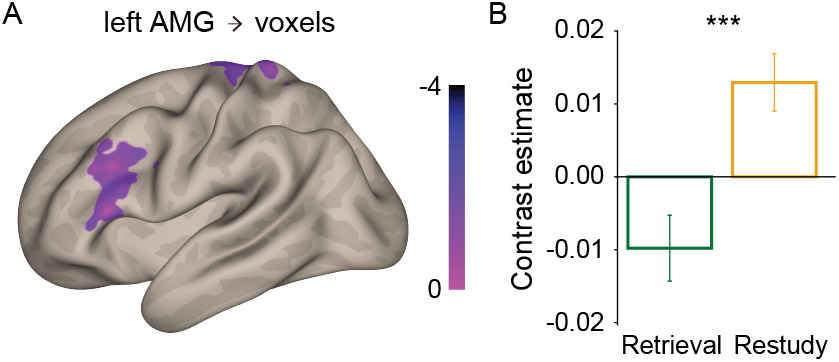
Tasked-modulated functional connectivity with the left AMG. The left AMG was used as a seed for gPPI analysis. Significant connectivity differences were observed in the left VLPFC (**(A)**), with negative coupling being observed in the Retrieval condition (**(B)**). Voxels shown at a peak threshold of p < 0.05 two-tailed, and a cluster threshold of p < 0.001, FWE-corrected. *** p < .001 (Two-tailed t test). Error bars indicate the standard errors of the mean.

### Relationships between selective retrieval and cognitive reappraisal

The foregoing findings demonstrate that even selective retrieval of content irrelevant to reinterpreting a negative scene (as is true with our unrelated neutral target words) is sufficient to regulate affective representations, indicating that the reappraisal itself is not necessary to achieve at least some emotion regulation benefits. These findings suggest that affective reappraisal regulates emotion, in part, because it engages selective retrieval to develop substitute thoughts and interpretations about a memory (Engen & Anderson, 2018). This hypothesis implies that reappraisal can be viewed as a specific form of thought substitution, a process known to involve selective retrieval and inhibitory control (Benoit & Anderson, 2012). If so, the brain regions recruited during selective retrieval should be related to those involved in thought substitution, and to those involved in reappraisal itself. To test these predictions, we assessed whether (a) thought substitution and reappraisal regions were engaged during our selective retrieval task and b) whether these regions showed hallmark characteristics associated with memory inhibition.

First, we tested whether selective retrieval recruited regions shared with thought substitution (TS), a strategy used to intentionally diminish the accessibility of negative content. Prior work by Benoit and Anderson (2012) identified two clusters within left VLPFC (i.e., left caudal PFC and mid-VLPFC) that are engaged when people try to prevent retrieval of a target memory, given a cue, by recalling distracting content to occupy awareness. Using these areas as ROIs for Thought Substitution in the current analysis, we found a substantial overlap between these regions and the left VLPFC clusters activated during selective Retrieval (Fig. 7A left). Indeed, when we restricted analysis of the current data to only those voxels within these a priori thought substitution ROIs, we found greater activation during Retrieval than Restudy (Fig. 7A right, t(27) = 2.55, p = .017, Cohen’s d = 0.48). These findings indicate that left PFC regions recruited during selective retrieval partially co-localize with those engaged during thought substitution, as would be expected, given that both manipulations encourage selective retrieval, though for distinct purposes (retrieval, versus self-distraction).

**Figure 7.**
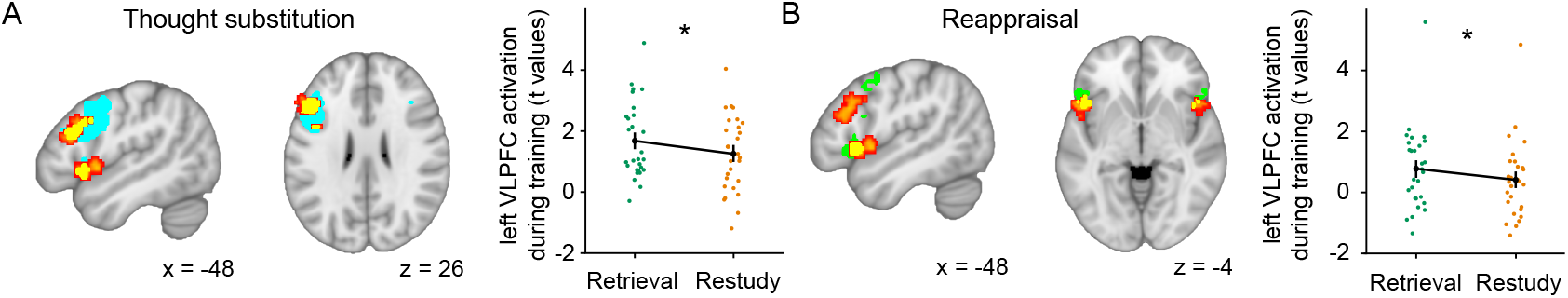
Relationships of prefrontal regions engaged by selective retrieval to those involved in thought substitution and reappraisal. **(A)** (left) Brain regions contributing to selective retrieval (red-orange) and intentional thought substitution (blue) from (Benoit & Anderson, 2012), with overlapping regions shown in yellow. (right) Activation in the left VLPFC region engaged by thought substitution showed greater activation during Retrieval than during Restudy. **(B)** (left) Brain regions contributing to selective retrieval (red-orange) and cognitive reappraisal (green) from (Buhle et al., 2014), with overlapping regions shown in yellow. (right) Activation in the left VLPFC engaged by cognitive reappraisal showed greater activation during Retrieval than during Restudy. * p < .05 (Two-tailed t test). Error bars indicate the standard errors of the mean.

Next, we compared the regions involved in selective retrieval and cognitive reappraisal. Previous research on reappraisal consistently identifies increased neural activity in the left VLPFC, which has been suggested to support the selection of appropriate reappraisals (Buhle et al., 2014). To test whether these regions are recruited during selective retrieval, we isolated the two left VLPFC ROIs from the Buhle et al. (2014) meta-analysis, one left ventral IFG and one left posterior MFG. Indeed, the reappraisal ROI in the left ventral IFG overlapped with a cluster identified in our whole-brain analysis comparing activation during selective Retrieval with Restudy, in the left M/IFG (Fig. 7B left). Again, when we restricted our analysis of the current data to the left ventral IFG reappraisal cluster, we found that Retrieval was accompanied by higher activation than was Restudy (Fig. 7B right, t(27) = 2.20, p = .037, Cohen’s d = 0.42). Thus, the prefrontal region recruited during selective retrieval partially co-localizes with brain regions consistently engaged by cognitive reappraisal, consistent with the possibility that mnemonic inhibition processes contribute to the affect regulation benefits of reappraisal.

## Discussion

Here we examined whether selective retrieval may constitute a previously unrecognized mechanism of emotion regulation, as predicted by the selective retrieval hypothesis of cognitive reappraisal. Abundant prior work has shown that attempting to recall a particular target item from long-term memory disrupts the retention of competing traces, via top-down inhibitory control mechanisms mediated by the VLPFC. Whether and how such selective retrieval might also alter affective responding to unpleasant competing content has remained unexplored, however. In the current study, we found that selective retrieval not only impairs episodic memory for competing traces, but also attenuates emotional responding to those competitors. The mnemonic effects of selective retrieval were seen in reduced memory for competing unpleasant scenes, the decreased activity in memory-related brain regions, and the disrupted neural representations in VTC; affective changes instead were reflected by decreased skin conductance response to competing scenes as well as reduced neural activation and disrupted neural representations in the amygdala unique to the unpleasant competitors. Taken together, these findings indicate that selective retrieval can disrupt the retention of unpleasant scenes and alter affective responding to them, possibly via parallel and distinct effects on mnemonic and affective regions.

Our results offer direct and multifaceted support for the inhibitory control account of RIF. Behaviorally, RIF exhibits several distinctive functional properties, including retrieval specificity and strength independence (Anderson & Hulbert, 2021; Anderson & Spellman, 1995), all consistently observed in our results. Neurologically, our findings establish a connection between emotion and memory regulation through selective retrieval via the left VLPFC, a region that resolves retrieval competition and adaptively disengages when retrieval competition is resolved (Badre & Wagner, 2007; Kuhl et al., 2007). However, the functional properties of forgetting only provide indirect support for inhibition (Wimber et al., 2015). To address this issue, we included a unique manipulation by introducing an emotional dimension exclusively to the competitors (i.e., the competitors were emotionally negative while the targets were emotionally neutral). Because retrieval of the neutral targets would be unlikely to activate emotion-related brain regions (Buchanan, 2007), changes in amygdala responses during retrieval should provide a clean measure of the impact of prefrontal control on the affective competitors. We observed below-baseline activation in the amygdala when competition from unpleasant competing scenes was being overcome, providing evidence that those competitors were actively inhibited. Consistent with these findings, our template-based multivariate imaging analysis revealed diminished representations of both affective and episodic contents for the unretrieved competitors in regions associated with emotion and memory, respectively. Collectively, these findings present evidence that selective retrieval induces inhibition of memory and emotion representations associated with competing memories.

Evidence for the suppression of mnemonic and affective representations arose in the VTC and the left amygdala respectively, in line with these regions’ roles in memory and emotion processing. However, the hippocampus displayed inconsistent evidence for memory suppression (e.g., reduced univariate activity, but no relationship of this to forgetting). This result, in fact, is consistent with previous research related to retrieval-induced forgetting, which often has not observed robust evidence for disrupted neural patterns in the hippocampus in RIF (Wimber et al., 2015). One candidate explanation for this is the concurrent presence of both the retrieval process (needed to recall the target item) and the suppression mechanism (needed to suppress the competitor) in the RIF paradigm, two requirements likely to place opposing demands on hippocampal activity that may counteract each other. Interestingly, we still observed a modest reduction in univariate signal in the hippocampus in the selective retrieval compared to the restudy condition. This discrepancy could be attributed to our use of distinct stimuli for retrieval (i.e., words) and suppression (i.e., pictures). A similar rationale may account for the observed decrease in activity within VTC. Whereas we did not consistently observe a relationship between hippocampal activity and forgetting, we established a connection between the retrieval-induced forgetting effect and alterations in both univariate and multivariate patterns in the VTC. This finding is particularly noteworthy as the VTC plays a crucial role in memory reinstatement (Wang et al., 2023; Xiao et al., 2017).

Reappraisal, recognized as an essential strategy for emotion regulation associated with wide-ranging improvements in psychological well-being, has found widespread application both in daily life and in clinical practice (Aldao et al., 2010). Nevertheless, the neural mechanisms contributing to the emotional benefits of reappraisal remain unclear. One critical reason is that the conventional cognitive reappraisal procedure, which involves positive reinterpretation of an existing event, does not enable one to discern whether the emotional benefits result from the suppression of negative content or depend on the presence of a positive framing of the stimulus. In our experimental design, we replaced reappraisal with the retrieval of neutral semantically unrelated words, effectively excluding the influence of positive reframing. Our findings of attenuated amygdala activity and emotional responding arising from selective neutral word retrieval thus indicate that suppressing negative memories alone may reduce negative emotional responses. This assertion was further supported by the observation that neither the SCR nor amygdala response changes were associated with the retrieval benefit on the neutral targets (ps > .05). Building on this, we delved deeper into whether the emotional benefits arose from the reduced accessibility of memories due to RIF or whether selective retrieval directly suppressed emotional responses independently of the forgetting effect. Supporting the latter perspective, we observed decreased amygdala activity to a level below that of the fixation baseline in the Retrieval condition. This decrease in amygdala activity cannot be solely attributed to reduced accessibility of upsetting event details, as a decrease in memory would, at most, eliminate emotional responses that otherwise would have been elicited by a retrieved competitor memory, but would not have reversed them. Furthermore, our investigation showed that the emotional benefits, as reflected in the decrease in SCR and inhibition of amygdala response, were unrelated to changes in memory accessibility of competing scenes.

The present findings support the possibility that inhibitory mechanisms recruited during selective retrieval may be a mechanism for reappraisal, consistent with the selective retrieval hypothesis of cognitive reappraisal (Engen & Anderson, 2018). Reappraisal and selective retrieval share conceptual similarities, with both involving an active retrieval process that ultimately diminishes the emotional responses to the unwanted event. Here we found that these similarities are more than conceptual: both tasks engage shared brain regions. Our findings confirmed, for example, that the left VLPFC regions of interest for selective retrieval practice, thought substitution (Benoit & Anderson, 2012), and reappraisal all share the feature that they all exhibit greater responsiveness to active retrieval than they do to a restudy task in which the same content is reprocessed, but without the need to overcome retrieval competition. Importantly, we observed that our selective retrieval task downregulates neural activation in the left amygdala, an effect that aligns with the emotion regulation consequences of reappraisal (Buhle et al., 2014; Etkin et al., 2015). However, further work is needed to firmly establish the role of selective retrieval in reappraisal’s affective benefits. First, if the inhibitory process supporting selective retrieval underlies reappraisal, the emotion regulation effects arising in the former may predict that observed in the latter. Similarly, enhancing inhibitory control ability should benefit reappraisal ability. Secondly, if reappraisal engages selective retrieval, reappraisal should influence episodic memory. This prediction derives from the impact of selective retrieval on activation of and representations in VTC, and their relationship to activation in left vLPFC. However, whereas our findings suggest that selective retrieval contributes on cognitive reappraisal, we do not discount the impact of other factors (e.g., positive framing, and executive control abilities aside from inhibition) on reappraisal’s effect.

Collectively, our study suggests evidence that inhibitory processes engaged by selective retrieval diminish emotional responses to competing memories via interactions between the left VLPFC and the amygdala. RIF has been proposed to be a reflection of mechanisms that are essential to adaptive forgetting that are present in humans and other mammalian brains (Bekinschtein et al., 2018; Storm & Jobe, 2012). Here, we provide further evidence that, in addition to its role in memory control, the inhibitory processes underlying RIF modulate individuals’ emotional responses to affective memories. Given that many emotional experiences in life derive from the emotional content of our memories, how well people regulate memories via inhibitory control may prove to be an important contributor to emotion regulation outcomes (Engen & Anderson, 2018; Rowlands et al., 2025).

## Materials and methods

### Participants

Thirty-six healthy college students participated in the behavioral experiment (15 males, mean age = 22.7 years) and thirty-two (8 males, mean age = 20.4 years) participated in the fMRI experiment. Four participants in the fMRI experiment were excluded from further analysis, three for falling asleep in the scanner and one because recall responses were not recorded; of the remaining 28 participants on whom skin conductance response data were collected, data recording problems made data from 5 participants unusable (i.e., data file corrupt, N = 2; physiological data not stored, N = 3). The sample size of the behavioral experiment was comparable to those in the field; the sample size of the fMRI experiment was determined based on a priori power analysis (power = 0.85, one-tail alpha = 0.05) of the RIF effect in the behavioral experiment. The study was approved by the human subject review committee at Peking University. Written consent was obtained from each subject before starting the experiment.

### Materials

The stimuli consisted of 48 object-scene pairs (competitor pairs) and 48 object-word pairs (target pairs). The object-scene pairs were selected from (Küpper et al., 2014). The scenes, which were originally taken from the International Affective Picture System and online sources, contained trauma themes such as physical and sexual assault, witnessing injuries and death, natural disasters, and serious accidents. The object pictures, cues for the scenes, were photographs of familiar objects that resembled an item embedded in its paired scene but that was not related to the scene’s central gist. The object-word pairs were composed by pairing each of the object pictures from the object-scene pairs with a two-character neutral Chinese word. Thus, each object cue was paired with both a scene and a word. The 48 object-scene/word pairs were split into three subsets of 16 pairs. For a given subject, each subset would be assigned to the Retrieval, Restudy, or Control conditions (as in (Ye et al., 2020)). Assignments of the three subsets to experimental conditions were counterbalanced across participants. A group of 16 scene pictures were further included in Experiment 2 to serve as a baseline condition for the emotional response test.

### Experimental Procedure

#### Experiment 1 (behavioral experiment)

To match the strength of initial encoding for pairs in different conditions, participants first studied all pairs through a drop-off/feedback cycle procedure. Participants studied the object-word (target) and object-scene (competitor) pair sets separately to discourage participants from integrating the word and the scene. After studying each object-word pair for 4 s (interstimulus interval, ISI = 1 s), participants were given test trials presenting the cue object alone for up to 4 s and were asked to judge whether they could retrieve the corresponding target word or not by pressing one of two keys. When a key was pressed or when the response window had expired, the target word appeared. Participants then judged whether they had retrieved the target word correctly or not. This procedure continued on pairs that had not been correctly recalled until 60% of the targets were indicated to be successfully recalled. After a short break, participants studied the object-scene pairs with the same procedure. Participants were informed that the cue objects in the two series were the same and were instructed to study the new pairs without thinking of the first set or integrating the three items (i.e., one common object cue, one word target, and one scene target) into a single memory. In the Retrieval/Restudy phase, object-word pairs from two of the three stimuli subsets were used for selective Retrieval and Restudy. During Retrieval trials, each cue object was presented for 4 s, and participants were instructed to retrieve the target word associated with the object. In Restudy trials, each cue object was presented together with its associated word such that participants did not need to retrieve the target, and participants were asked to repeat the object-word pair to themselves. Notably, both Retrieval and Restudy were performed on object-word pairs. The selective Retrieval or Restudy process repeated eight times for each object-word pair (over four blocks). The order of trials for different conditions was randomized with the restriction that no more than three consecutive trials belonged to the same condition. Following the Retrieval/Restudy phase, the aftereffects of Retrieval and Restudy were examined via a cued-recall test first on the object-scene pairs and then on the object-word pairs, in separate testing blocks. For the object-scene recall test, cue objects from all three subsets were presented on the screen individually, each for 15 s (ISI = 3 s), and participants were asked to describe the associated scene images in as much detail as possible. Scoring of identification success was made based on the criterion in (Küpper et al., 2014). Specifically, a description of the target scene was scored as correct if it included enough detail for an independent rater to identify the scene. Following our previous procedure (Zhu & Wang, 2021; Zhu et al., 2022), the description was scored by one trained coder who was blind to the conditions to maintain consistency. An independent coder was invited for a subsample of 10 participants, which showed a high interrater agreement (r = 1). The cued-recall test for object-word pairs followed, in which cues from all three subsets were presented on the screen individually, each for 3 s, and participants reported the paired word. Verbal responses were recorded.

#### Experiment 2 (fMRI experiment)

The fMRI experiment adopted the same procedure as in Experiment 1, with two additional phases: a re-exposure phase and an emotional response test. The re-exposure phase was conducted inside the scanner, right before the Retrieval/Restudy phase, for the purpose of obtaining a perceptual template of each competitor scene image. During re-exposure, scene images from the object-scene pairs appeared on the screen one by one, each for 4 s. Participants were asked to look at the content of the images. To avoid memory differences in the three experimental conditions caused by the re-exposure, all 48 scene images from the three subsets appeared during this phase. Each image was re-exposed twice, in two separate runs. The ISI was generated using OptSeq2, with a jitter range of 0-4 s. The Retrieval/Restudy phase followed the re-exposure phase and was completed in 4 or 8 runs inside the scanner. Instead of using a fixed ISI as in Experiment 1, the ISIs for Retrieval/Restudy were generated using OptSeq2, with a jitter range of 0-6 s. The total duration of each re-exposure run was 350 s and that of each Retrieval/Restudy run was 238 s. In addition, an emotional response test was included to measure participants’ subjective and objective emotional responses to the competitor scene images. To measure subjective emotional responses, we used a Valence and Arousal rating task. Each image appeared for 10 s, with 7-point Likert scales for Valence and Arousal appearing sequentially below the image each for 5 s. Participants rated their subjective feelings about the image using the scales. For the psychophysiological indices of emotional responses, participants’ skin conductance responses were recorded, during the Valence/Arousal rating task. To control for Valence/Arousal differences in stimuli across conditions, these emotional tests were given twice, once at the beginning and once at the end of the experiment. The effect of Retrieval/Restudy on the emotional response to the competitive memories is thus assessed by examining changes in Valence/Arousal ratings or in SCR responses after compared to before the Retrieval/Restudy task. In addition to the 48 scene images from the three experimental conditions (i.e., Retrieval, Restudy, and Control), a set of 16 scene images that were not presented in the other phases of the experiment were presented, serving as a Baseline to show pure effect of stimuli repetition. Images from different conditions were presented in a random order.

### Skin Conductance Response (SCR) Analysis

#### SCR recordings and preprocessing

Skin conductance was recorded on the volar surface of the distal phalanx of the index and middle fingers of the nondominant hand (the hand that was not used for key pressing during Valance/Arousal rating) using 8 mm Ag/AgCl cup electrodes (EL258, Biopac Systems, Goleta CA, USA) and 0.5%-NaCl electrode paste. Due to technical issues, we only obtained SCR data from 23 valid participants. Skin conductance was recorded at a constant voltage of and sampled at 1000 Hz. Following current recommendations (Bach et al., 2013), SCR data was imported, trimmed and preprocessed using PsPM (version 4.0.2). The data was filtered using a unidirectional first order Butterworth high pass filter with cut off frequency 0.05 Hz. This was done to account for the ‘tonic’ changes in skin conductance, over the course of the experimental sessions (Lim et al., 1997).

#### SCR data analysis

After preprocessing, the time series for each session, for each participant, was extracted in PsPM. The marker channel was imported to PsPM along with the skin conductance data channel. The counterbalancing order for the three experimental conditions was also taken into account, in order to correctly classify each trial into the corresponding experimental condition it belonged to. Then, the extracted time-series was averaged over all the trials in a particular condition, to get an average time-series for each condition. This was done for both Pre- and the Post-experiment sessions. We used the first one second of stimulus presentation, across conditions, to compute the skin conductance level (SCL) for every participant, for each of the two sessions. The first one second after stimulus presentation is widely acknowledged to be the “latency” period for eventrelated evoked skin conductance responses (SCRs) (Bach et al., 2010; Braithwaite et al., 2013). Hence, this can be taken to reflect a subject-specific baseline SCL, and was subtracted from the rest of the extracted time-series. The rest of the time series was deemed to belong to a canonical evoked SCR, and the length of each SCR was the length of the stimulus presentation minus one second (so 9 seconds in total). The area under the curve (AUC) was calculated for each condition, each participant, separately for each of the two sessions. This way, we could quantify the effect of the Retrieval/Restudy on the SCR evoked for the different scenes, for each participant. This is the same approach shown earlier for spontaneous skin conductance fluctuations (Bach et al., 2010), and has been subsequently used to model event-related evoked SCRs (Harrington et al., 2021). The data were then entered into group level comparisons and correlation analyses.

### fMRI Analysis

#### MRI acquisition

Whole-brain MRI data were collected on a 3.0T Siemens Prisma MRI scanner at the Center for MRI Research at Peking University. A high-resolution simultaneous multislice EPI sequence was used for functional scanning (62 slices, repetition time (TR) = 2000 ms, echo time (TE) = 30 ms, flip angle = 90°, field-of-view (FOV) = 224 × 224 mm, matrix = 112 × 112, slice thickness = 2 mm, slice gap = 0.3 mm, GRAPPA factor = 2, multi-band acceleration factor = 2). A high-resolution structural image using a 3D T1-weighted MPRAGE sequence was acquired to cover the whole brain (TR = 2530 ms, TE = 2.98 ms, flip angle = 7°, FOV = 256 × 256 mm, matrix = 256 × 256, slice thickness = 1 mm, GRAPPA factor = 2).

#### Image preprocessing

MRI data were preprocessed using FMRIPREP V1.5.4. Functional data were slice time corrected using AFNI v16.2.07, motion-corrected using FSL’s MCFLIRT, and registered to the T1 image using a boundary-based registration with nine degrees of freedom. The T1 volumes were skull-stripped using antsBrainExtraction.sh (OASIS template). Cortical surfaces were reconstructed using FreeSurfer v6.0.1. The T1 volumes were normalized to the ICBM 152 Nonlinear Asymmetrical template (version 2009c) through nonlinear registration with the ANTs v2.1.0. For the univariate analysis, data were spatially smoothed with a 4 mm full-width-at-half-maximum (FWHM) Gaussian kernel using FSL’s SUSAN, filtered in the temporal domain using a nonlinear high-pass filter with a 100 s cutoff, and normalized to MNI template space. For the representational similarity analysis, data were spatially smoothed with a 1.6 mm FWHM Gaussian kernel, aligned to subjects’ T1 image, and kept at its native resolution.

#### Univariate analysis

We used the general linear model within the FILM (version 6.00) module of FSL to model the two types of trials: The Retrieval and Restudy trials. The regressors were convolved with a double gamma hemodynamic response function (HRF). Six movement parameters and the framewise displacement (FD) were modeled as confound regressors. Additional censor regressors were included for each volume with an FD greater than 0.3 mm. Each run was modeled separately in the first-level analysis. Cross-run averages for each contrast image were created for each subject with a fixed-effects model. These contrast images were then entered into the group level random-effect statistical tests. Group images were thresholded using a cluster-size test, with a threshold of z > 2.3 and a cluster probability of p < 0.05, combined with random field theory for inference.

#### Definition of Regions of interest (ROIs)

Four ROIs (the left middle/inferior frontal gyrus, the left hippocampus, the left amygdala, and VTC) were selected based on their associations with episodic retrieval and emotional memory processing. They were defined using Freesurfer Destrieux atlas (the following label numbers refer to Simple-surface-labels2009.txt). The left middle/inferior frontal gyrus ROI was comprised of regions 12 - 15, 39, 40, 53, 54, and 63. The left HPC ROI was comprised of region 17. The left AMG ROI was comprised of region 18. The VTC ROI was comprised of regions 21, 23, 51, 52, 61, and 62. All ROIs were originally defined in MNI space, and were realigned to each subject’s native anatomical space for the representational similarity analysis. For ROI-level univariate analysis, we extracted the t-statistics for voxels within the ROI from the GLM analysis on the Retrieval and Restudy conditions and calculated the averaged t-statistics across voxels for each condition.

#### Single-trial response estimate

The GLM models were separately created for each of the 32 Retrieval and Restudy trials in each run to estimate the single-trial response. A least-square single method was used for each trial, where the given trial was modeled as a separate regressor and all the remaining trials were modeled as another regressor. The trial was modeled at its presentation time, convolved with a double gamma HRF. The same confounding regressors were included as in the univariate analysis. The t-statistics were used for representation similarity analysis to increase the reliability by noise normalization (Walther et al., 2016).

#### Representational similarity analysis (RSA)

RSA was used to determine the degree of competitive memory reactivation during Retrieval and Restudy. Each trial’s t-statistic values from the single-trial response estimations within each ROI were extracted. For each scene image in each run, Pearson correlation was performed between the averaged activation pattern during re-exposure (i.e., the perceptual template) and the activation pattern during Retrieval or Restudy of its competing object-word pair. The correlation thus represented the reactivation level of the competing scene image during Retrieval or Restudy. The Fisher Z-transformed correlations were then grouped and averaged based on the training repetitions and the training conditions for further statistical analysis. To explore the representation changes along with Retrieval practice in the amygdala, we split the eight Retrieval/Restudy runs into four sections (two runs per section) and computed a linear fit of the representation similarities across the four sections using the linear fit function in MATLAB. Paired t-tests were used for group-level analysis.

#### Functional connectivity analysis

PsychoPhysiological Interaction (PPI) measures the temporal relation between a seed region and a target region after partialling out the common driving influence of task activity on both regions, thereby enabling inference regarding condition-specific functional integration. We used a generalized form of context-dependent PPI (gPPI) analysis (McLaren et al., 2012) implemented in the functional connectivity toolbox Conn (Whitfield-Gabrieli & Nieto-Castanon, 2012). The gPPI method has the advantage of spanning the entire experimental space over the standard PPI, allowing explorations of connectivity in individual conditions within the experiment. At the individual participant level, we included: (1) a psychological variable representing the two types of conditions (i.e., Retrieval and Restudy), (2) one physiological variable (i.e., the time course in the seed region), and (3) a PPI term (i.e., the product of the first two regressors). Here we used a combined ROIF of the left hippocampus and the left amygdala as a seed region and explored its functional connectivity with regions spanning the whole brain. The gPPI statistics were thresholded by a voxel threshold of p < 0.05 two sided, corrected at the cluster-level using a family-wise error rate of p < 0.001.

### Correlation on Normalized Data

Behavioral and neural measures were z-normalized within that participant’s counterbalancing group before being submitted for correlation analysis. Z-normalizing within each item counterbalancing group controls for differences in the memorability of items in each counterbalancing set by quantifying how unusual a participant’s behavioral or neural response is with respect to a homogenous group of participants receiving precisely the same items in the same conditions (Anderson et al., 2004; Nardo & Anderson, 2024).

